# Tumor Immunity Microenvironment-based classifications of bladder cancer for enhancing cancer immunotherapy

**DOI:** 10.1101/2020.08.22.262543

**Authors:** Jialin Meng, Xiaofan Lu, Yujie Zhou, Meng Zhang, Jun Zhou, Zongyao Hao, Yinan Du, Fangrong Yan, Chaozhao Liang

## Abstract

**Background:** Bladder cancer is composed by a mass of heterogenetic characteristics, immunotherapy is a potential way to save the life of bladder cancer patients, but only benefit to about 20% patients.

**Methods and materials:** A total of 4003 bladder cancer patients from 19 cohorts was enrolled in this study, collecting the clinical information and mRNA expression profile. The unsupervised non-negative matrix factorization (NMF) and nearest template prediction (NTP) algorithm was used to divide the patients to immune activated, immune exhausted and non-immune class. Verified gene sets of signatures were used to illustrate the characteristic of immunophenotypes. Clinical and genetic features were compared in different immunophenotypes.

**Results:** We identified the immune class and non-immune classes in from TCGA-BLCA cohort. The 150 top different expression genes between these two classes was extracted as the input profile for the reappearing of the classification in the other 19 cohorts. As to the activated and exhausted subgroups, a stromal activation signature was conducted by NTP algorithm. Patients in the immune classes shown the highly enriched signatures of immunocytes, while the exhausted subgroup also shown an increased signature of TITR, WNT/TGF-β, TGF-β1 activated, and C-ECM signatures. Patients in the immune activated shown a lower CNA burden, better overall survival, and favorable response to anti-PD-1 therapy.

**Conclusion:** We defined and validated a novel classifier among the 4003 bladder cancer patients. Anti-PD-1 immunotherapy could benefit more for the patients belong to immune activated subgroup, while ICB therapy plus TGF-β inhibitor or EP300 inhibitor might be more effectiveness for patients in immune exhausted subgroup.

## INTRODUCTION

Bladder cancer is the 10^th^ most frequent tumor globally, of which with a high rate of recurrence^1^. There are about 550 thousand new cases and 200 thousand specific deaths caused by bladder cancer each year. The incidence rate of bladder cancer variable around the world, with the highest rate in Southern Europe, and lowest rate at Middle Africa^2^. In United States, bladder cancer ranges the 6^th^ common tumor, with about 81 thousand new cases and 18 thousand new deaths at 2020^3^. Most patients diagnosed at the early stage of bladder cancer, also known as the non-muscle-invasive bladder cancer (NMIBC), which only processes away from the muscle layer and could be removed easily through the transurethral resection (TUR), or plus with the intravesical therapy with Bacillus Calmette-Guérin (BCG) or other chemotherapeutic medicine ^4-6^. Recurrence is extremely common in NMIBC, about 70% patients will suffer from the health burden of bladder cancer again within 10 years, and one thirds of them step into advanced stage, also called muscle-invasive bladder cancer (MIBC)^7^. The standard care of MIBC is radical cystectomy with or without neoadjuvant chemotherapy or chemoradiation. And even after the treatment, almost 50% MIBC patients will recurrent and death from the metastatic stage within 3 years^8^.

In the tumor mass, the normal cells, blood vessels, and cytokines which surround and support the alive of tumor cells are composes of the tumor microenvironmnt (TIME). The cross talk existed between the tumor and the TIME, the tumor cells could alter the TIME, and the TIME could also promote the growth and spread of the tumors^9^. Several studies investigated the diagnosis markers, prognostic signatures or therapeutic targets for malignancy tumors based on the TIME, as well as in bladder cancer. BCG is the earliest immune therapy approved in bladder cancer treatment, which could stimulate an immunologic reaction inducing a proinflammatory cytokine and direct cell-to-cell cytotoxicity^10^. BCG is still the standard therapy for NMIBC, which reflect that bladder cancer patients could benefit from immunotherapy. The development of the blockade of immune checkpoints also applied in the treatment of bladder cancer. Two PD-1 inhibitors (pembrolizumab and nivolumab) and three PD-L1 inhibitors (atezolizumab, durvalumab, and avelumab) were approved by the FDA for the treatment of bladder cancer (https://www.accessdata.fda.gov/scripts/cder/daf). In the IMvigor210 clinical study, atezolizumab was used to block the PD-L1, the objective response rate (ORR) for IMvigor210 cohort 1 is only 23%, 15% in the cohort 2^11,12^. The ORR for Nivolumab and durvalumab is similar, from 17% to 24.4%^13-16^. Therefore, a full understanding of the immunophenotypes to bladder cancer is essential, which could as the guidance to choose the patients to receive the appropriate immunotherapy.

We enrolled 4003 bladder cancer patients from 20 independent cohorts. Non-negative matrix factorization (NMF) algorithm and nearest template prediction (NTP) was applied to distinguish the immunophenotypes of bladder cancer patients in training cohort of TCGA-BLCA, as well as validated in the other 19 cohorts. The novel definition of immunophenotypes could provide illuminations for the immunotherapy of bladder cancer patients.

## MATERIALS AND METHODS

### Bladder cancer patient cohorts

4003 bladder cancer patients were registered in the current study, with the gene expression profiles, clinicopathological information and overall survival data (**Figure 1**). For TCGA-BLCA cohort, we obtained the level 3 gene expression profile of 408 patients from TCGA Data Portal (https://tcga-data.nci.nih.gov/tcga), only genes expressed in at least 50% of the samples were retained for analyses. For the further external validation cohorts, GSE32894, GSE83586, GSE87304, GSE128702, GSE13507, GSE129871, GSE120736, GSE39016, GSE128701, GSE124035, GSE86411, GSE48276, GSE31684, GSE134292, GSE69795, the gene expression profile were collected from Gene Expression Omnibus (http://www.ncbi.nlm.nih.gov/geo/). For E-MTAB-4321 and E-MTAB-1803 cohorts, the gene expression profile were downloaded from ArrayExpress (https://www.ebi.ac.uk/arrayexpress/). For IMvigor210 cohort, the gene expression profile was obtained from IMvigor210CoreBiologies, a data package for the R statistical computing environment. Detailed information of these datasets was displayed in **Table S1**.

**Figure 1.**
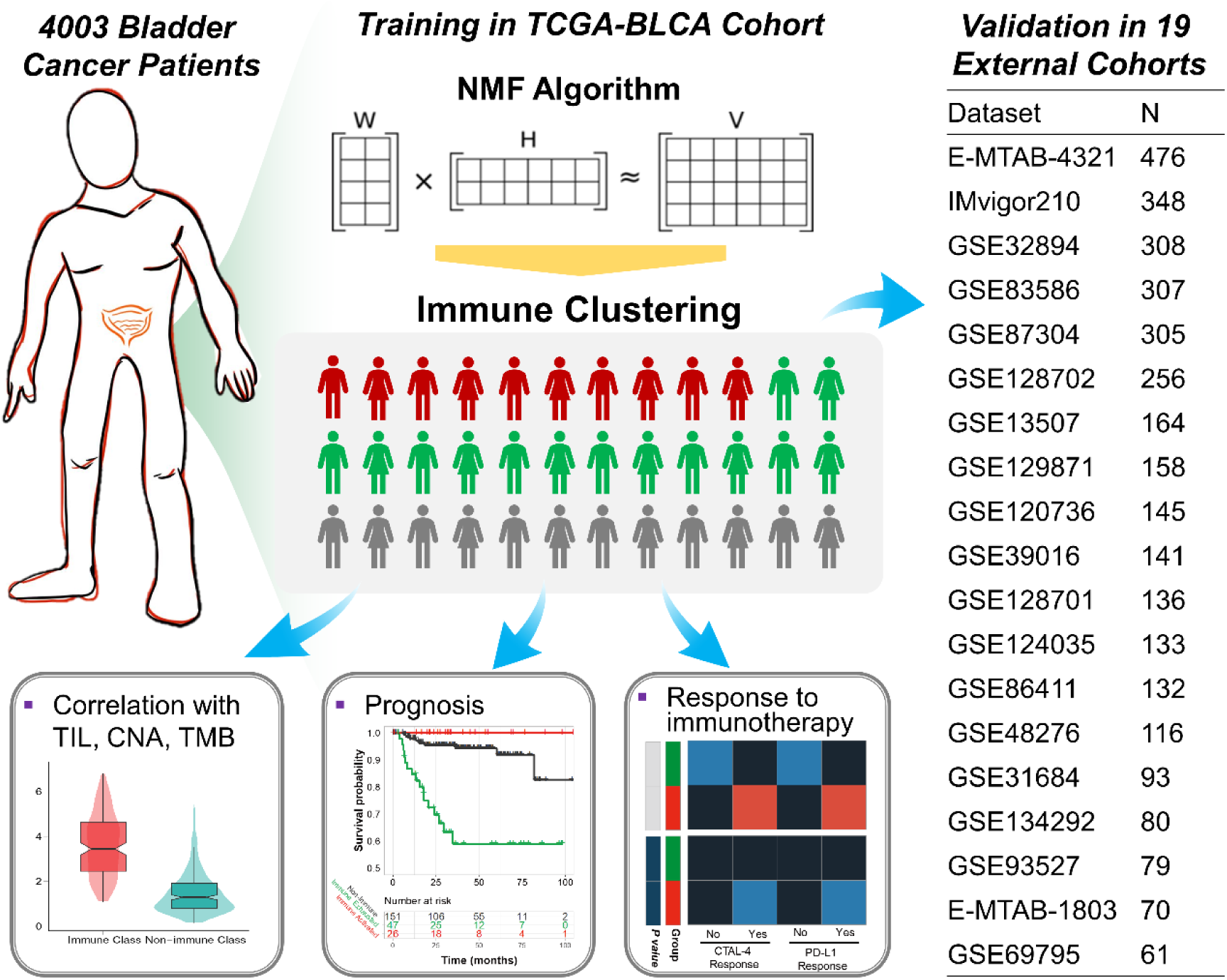
The flow chart demonstrates the summary of performed analysis in this study. A total of 4,003 bladder cancer patients from 20 cohorts with the mRNA expression profile were enrolled for the analysis, with non-negative matrix factorization algorithm and nearest template prediction, three immunophenotypes were generated in TCGA-BLCA cohort, and validated in 19 external cohorts. The molecular characteristic, prognosis and response to immunotherapy are difference in the three subtypes. NMF, non-negative matrix factorization; TCGA-BLCA, The Cancer Genome Atlas-prostate adenocarcinoma bladder cancer; TIL, tumor-infiltrating lymphocytes; CNA, copy number alteration; TMB, tumor mutation burden.

### Bioinformatic analyses

The mRNA expression profile of 408 bladder cancer patients from TCGA-BLCA cohort were microdissected by the unsupervised non-negative matrix factorization (NMF) algorithm^17^. Immune module was selected with the gathering of patients with the high immune enrichment score calculated by the the single-sample gene set enrichment analysis (ssGSEA) as described previously^18^. Top 150 exemplar genes with the highest weight in the immune module was extracted as the key genes to dichotomize the immune and non-immune classes, which further modified by the multidimensional scaling random forest method^19^. Immune activated and exhausted subgroups were recognized by the stromal activation signature with nearest template prediction^20^. To depict the characteristics of these three immunophenotypes, several immune associated signatures were manually collected and the score of each signature for each patient were generated via ssGSEA (**Table S2**). The different genetic types among immune and non-immune classes were evaluated, including tumor-infiltrating lymphocytes (TILs) abundance, Programmed death-ligand 1 (PD-L1) expression, Copy number alterations (CNA), tumor mutation burden (TMB), neoantigens and mutant genes. To reappearing the immunophenotypes we generated, the expression profile of the top 150 differentially expressed genes (DEGs), which increased in immune class than non-immune class, were used to dichotomize the immune classes in validation cohorts with the NMFConsensus method, immune class divided into activated and exhausted subgroups subsequently. Details about the enrolled cohosts, as well as the specific method of each step were provided in the **Supplementary Materials and Methods**.

## RESULTS

### Identify the immune module and derivate immune class of bladder cancer

We performed the virtual microdissection with the NMF algorithm, the mRNA expression profile of 408 bladder cancer patients from TCGA-BLCA cohort was analyzed as the training cohort. To obtain the robust immune module, we pre-set the numbers of module as five to 10, respectively, finally, when the total modules is nine, the first module strongly enriched the patients with a highly immune enrichment score (IES), of which defined as the immune module (**Figure 2A**). The top 150 weighted genes in the immune module were defined as the exemplar genes which could inflect the characteristics of the immune module (**Table S3**). These genes were involved in the signaling pathways of T cell activation, antigen processing and presentation, B cell activation based on the analysis among ontology biological process, and associated with the activation of the pathways related with Th1/Th2 cell differentiation, T cell receptor signaling pathway, B cell receptor signaling pathway, and PD-L1 expression/PD-1 checkpoint pathway (all, *P* < 0.05, **Table S4**). Subsequently, we re-defined the total 408 bladder patients to immune enriched or non-immune enriched groups by the consensus clustering based on the 150 exemplar genes (**Figure 2B**), what’s more, the multidimensional scaling (MDS) random forest was further employed to define a more precise classify of immune and non-immune classes (**Figure 2C**). In **Figure 2D**, the distribution and association of the 408 bladder cancer patients among NMF modules, immune module weight, exemplar gene clustering, final immune classes and immune enrichment score was shown.

**Figure 2.**
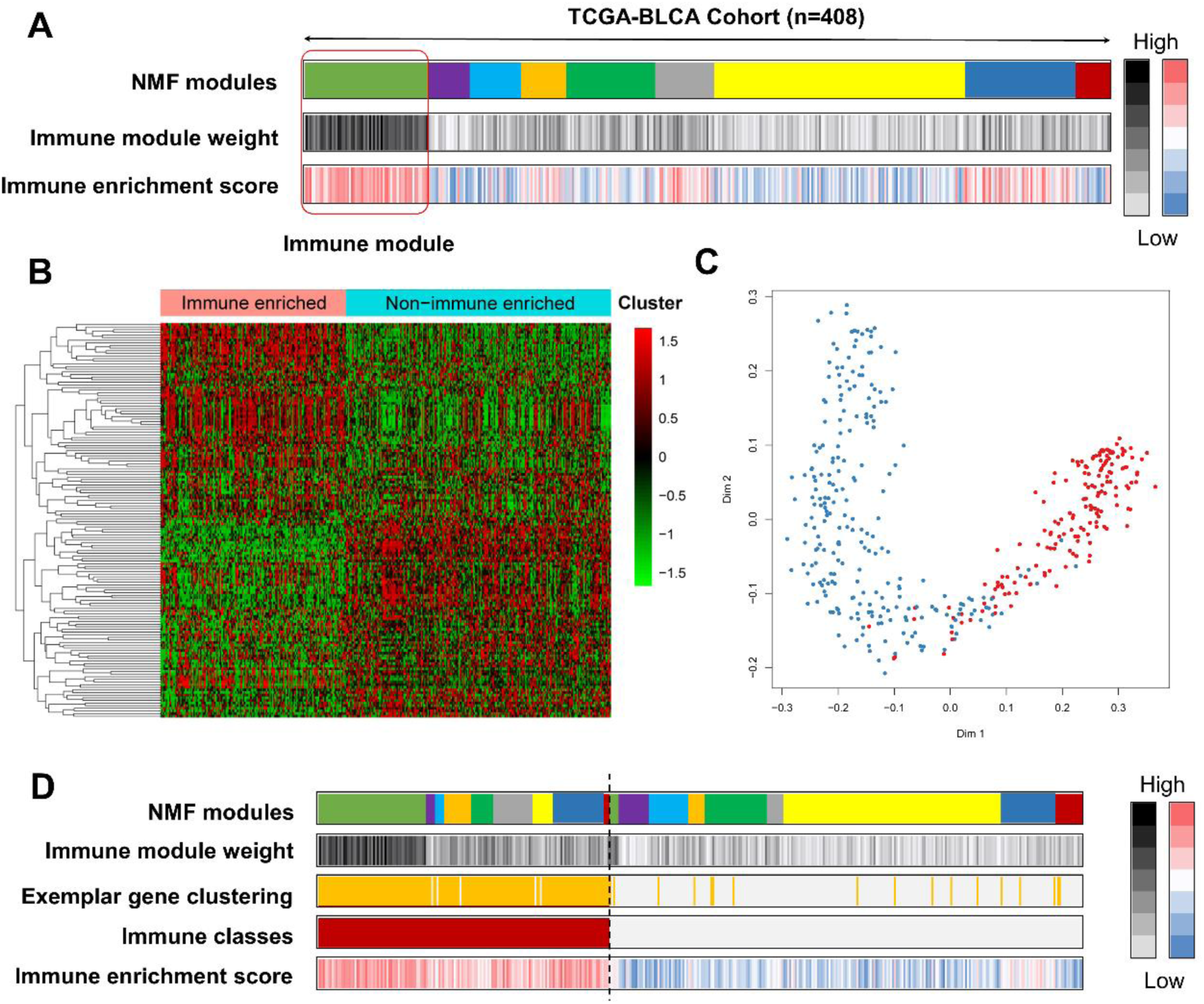
Recognition the immune classes by non-negative matrix factorization (NMF) algorithm. (A) Nine modules generated from the NMF algorithm, patients with high immune enrichment score gathered in the immune module; (B) Heatmap showing the top 150 exemplar genes expression among immune enriched and non-immune enriched clusters, divided by consensus clustering; (C) the multidimensional scaling (MDS) random forest further modified the clusters to immune and non-immune classes; (D) The distribution of patients in different NMF modules, immune module weight, exemplar gene clustering, final immune classes and immune enrichment score.

Several immune associated signatures (**Table S2**) were collected to help us to confirm the classification of immune or non-immune classes, the enrichment score of each signature among each patient were conducted by ssGSEA. We observed the increased enrichment of T cells (reflected by the signatures of 13 T-cell signature, T cells, CD8+ T cells, T. NK. Metagene), B cells (reflected by the signatures of B-cell cluster, B.P. meta), macrophages, tertiary lymphoid structure (TLS), cytolytic activity score (CYT) and IFN signatures (all, *P* < 0.05, **Figure 3**). We also analyzed the activated KEGG signaling pathways by GSEA, we revealed that the immune cell pathways (including T cell, B cell, natural killer cell and leukocyte associated pathways), immune response pathways (including chemokine signaling pathways, antigen processing presentation, cell adhesion molecules cams and complement coagulation cascades), proinflammatory pathways (including FC-Epsilon-RI, NOD like receptor and FC gamma R mediated phagocytosis pathways) were all active in the immune class (**Figure S1**). Taken together the results from **Figure 2, Figure 3 up panel, Figure S1**, and **Table S3-S4**, we microdissected the immune and non-immune classes in TCGA-BLCA cohort, activated immune associated signatures and signaling pathways were observed in the immune class.

**Figure 3.**
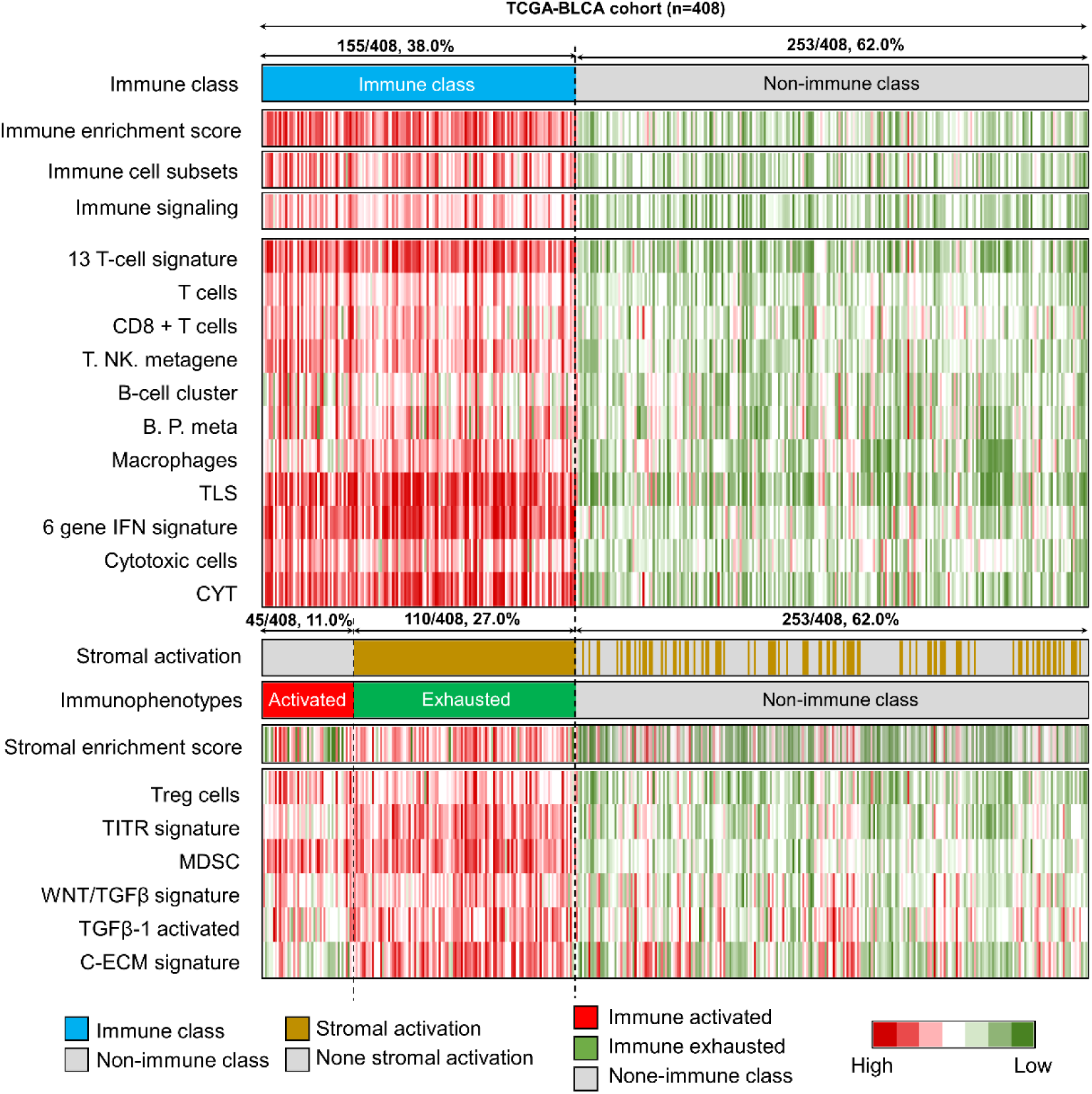
The diverse immune characteristics of non-immune class, immune activated subgroup, and immune exhausted subgroup. Immune class (253/408, 62.0%) and non-immune class (155/408, 38.0%) were distinguished by the consensus clustering and multidimensional scaling random forest based on the 150 exemplar genes obtained from the non-negative matrix factorization algorithm generated immune module; The immune activated subgroup (45/408, 11.0%) and immune exhausted subgroup (110/408, 27.0%) further divided by the stromal activation signature with nearest template prediction analysis. The high and low gene set enrichment scores are displayed with red and green, respectively. The details of these immune associated signatures listed in Table S2. TCGA-BLCA, The Cancer Genome Atlas-bladder cancer; CYT, cytolytic activity score; TITR, tumor-infiltrating Tregs; MDSC, myeloid-derived suppressor cell; TLS, tertiary lymphoid structure; C-ECM, cancer-associated extracellular matrix.

### Tumor immune microenvironment to immunophenotypes distinguished by activation of stromal cells

Fibroblasts, mesenchymal stromal cells (MSCs), and extracellular matrix (ECM) are the key components to compose the tumor stroma, support and connective the tumor cells^21^. Especially at the late stage of tumors, the genetic and epigenetic alterations of the tumor cells were driven by the activated stroma components^22^. MSCs act as the inherently regulators of tumor, which could secreta the inhibiting soluble factors and alter the cell surface markers to suppress the immune microenvironment, the inhibition of T-cell proliferation and induction of T regulatory cells (Tregs) were all affected by the regulation of PD-L1 by MSCs^23,24^. MSCs could handle the function of suppress immune process through decreasing the expression of pro-inflammatory factors, including IFN-γ, TNF-α and IL-1β, or promoting the expression of type 2 factors, IL-10 and IL13^25,26 27^. For this reason, the previously defined stromal activated signature was employed to further divide the immune class to immune activated and exhausted immunophenotypes, which could reflect the immune response status. A total of 11.0% (45/408) bladder cancer patients in TCGA-BLCA cohort was defined with the activated immunophenotype and inactivated stromal phenotype, belong to immune activated subgroup, while the other 27.0% (110/408) patients belong to immune exhausted subgroup, with the activated stromal phenotype (**Figure 3**). The ECM cytokines (C-ECM) secreted by the fibroblasts could recruit the immunosuppressive cells, the TGF-β is an accepted immunosuppressor in the immune microenvironment, as well as the Treg cells and MDSC cells could reflect the immune exhausted status in TME^28-31^. These signatures were evaluated by ssGSEA analysis, and we revealed that the TITR, WNT/TGF-β, TGF-β1 activated, and C-ECM signatures were higher in the immune exhausted subgroup than activated subgroup (all, *P* < 0.05, **Figure 3, Figure S2**). TIM-3 and LAG3 are reported associated with the immune exhausted status^32,33^, we also generated the similar results in the immune activated and immune exhausted subgroups, increased TIM-3 (*P =* 0.008) and LAG3 (*P =* 0.218) were observed in the immune exhausted subgroup (**Figure S2**). Based on the results from **Figure 3 down panel** and **Figure S2**, we separated the immune class into immune activated and immune exhausted subgroups, stromal enrichment score, TITR, MDSC and WNT/TGF-β signatures increased in the immune exhausted subgroup, and also validated by the immune exhausted markers, TIM-3 and LAG3.

### The heterogeneity of genetic phenotypes among immune and non-immune classes

To confirm the infiltration of immunocytes among the immune and non-immune classes distinguished by the mRNA expression profile of the exemplar genes, we compared the tumor-infiltrating lymphocytes (TIL) abundance of the 408 bladder cancer patients, which was anteriorly estimated by the Hematoxylin-eosin staining (H&E) stained whole-slide images of TCGA samples^34^. We obtained the result that the TIL abundance is higher in the immune class than non-immune class (*P <* 0.001, **Figure 4A**), consistent with the definition of these two groups. What’s more, we also observed the high expressed PD-L1 level in the immune class than non-immune class (*P <* 0.001, **Figure 4B**). The gene copy number alteration (CNA), tumor mutant burden (TMB), and neoantigens was reported have the crosstalk with tumor immune activation. Patients in the non-immune class shown an increased level of deletion in both arm and focal level (*P*_Arm-del_ < 0.001, *P*_Focal-del_ = 0.007), but not the CNA amplification (*P*_Arm-Amp_ = 0.733, and *P*_Focal-Amp_ = 0.065) (**Figure 4C**), which reflected the positive association of immune infiltration and gene CNA deletion. With the online tool of TIMER, we double confirmed the association between immune infiltration and gene CNA deletion, the deep deletion and arm-level deletion of PD-1, PD-L1 and CTLA4, the three major immune checkpoints, linked with the decreased immunocytes infiltration, especially for CD4+ T cell, Neutrophil, and dendritic cell (**Figure S3)**.

**Figure 4.**
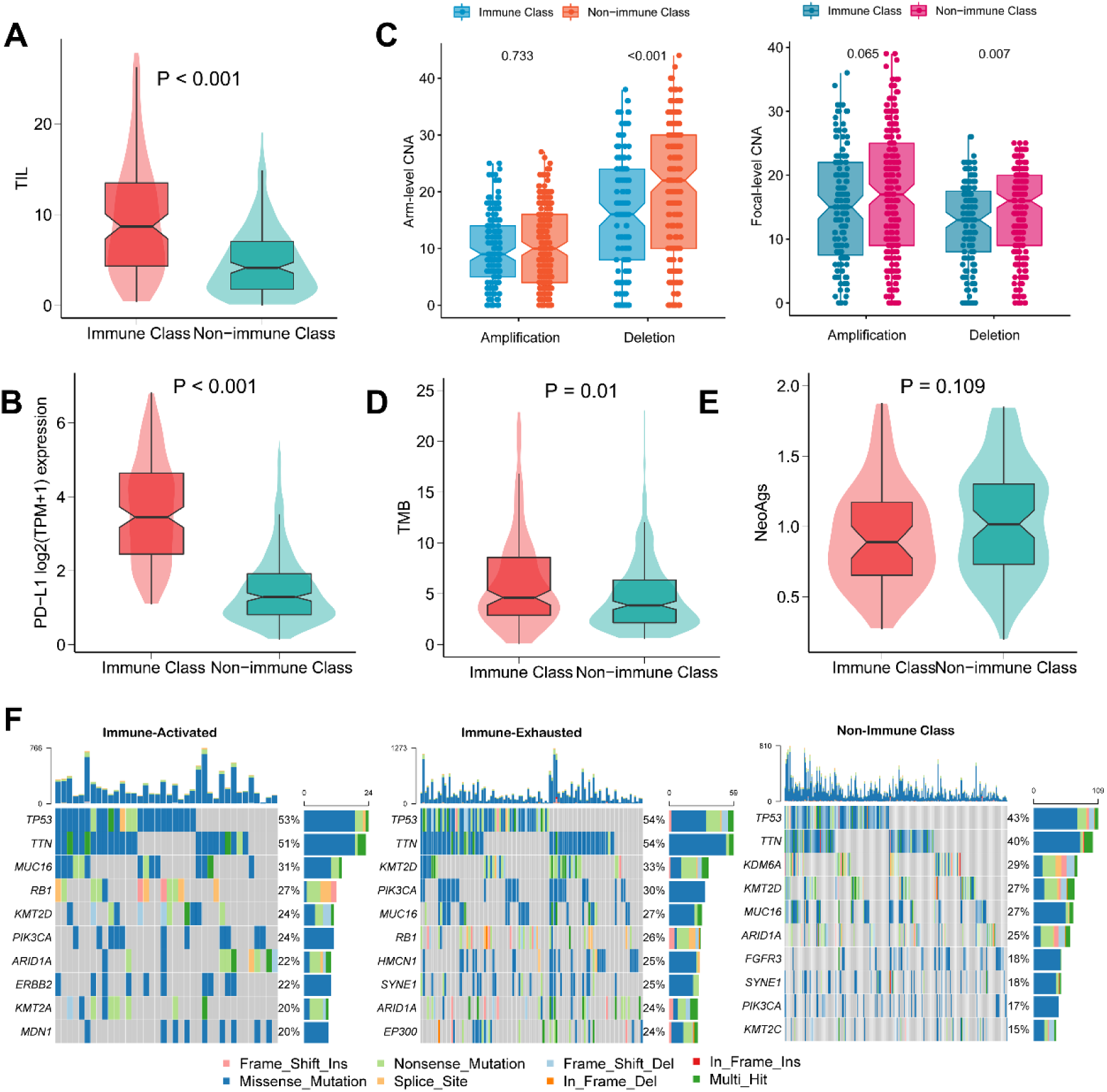
The heterogeneity of genetic phenotypes of non-immune class, immune activated subgroup, and immune exhausted subgroup. (A) Difference of tumor-infiltrating lymphocytes abundance; (B) Difference of PD-L1 mRNA expression level; (C) Difference of gene copy number alterations, including amplification and deletion, among arm-level and focal level; (D) Difference of tumor mutation burden; (E) Difference of neoantigens; (F) Different distribution of mutant genes in three immunophenotypes.

The TMB in immune class is higher than that in non-immune class (*P =* 0.01, **Figure 4D**), while the neoantigens level shown no difference (*P =* 0.109, **Figure 4E**). We further compared the specific gene mutations in the immune subgroups (**Figure 4F**). The mutation of TP53 (53.5% vs. 43.1%, *P =* 0.051), TTN (52.9% vs. 39.5%, *P =* 0.011), PIK3CA (28.0% vs. 17.0%, *P =* 0.007) and RB1 (26.0% vs. 13.0%, *P <* 0.001) appeared more in the immune class than non-immune class (**Figure 5A**). While ERBB2 (*P =* 0.035), KMT2A (*P =* 0.013), PKHD1 (*P =* 0.007) and MDN1 (*P =* 0.015) similar to be the specific mutations of immune activated subgroup(**Figure 5B**), and EP300 (*P =* 0.020), HMCN1 (*P =* 0.014), AKAP9 (*P =* 0.003) and MACF1 (*P =* 0.016) mutant patients enriched more in immune exhausted subgroups(**Figure 5C**). Taken together the results from **Figure 4, Figure 5**, and **Figure S3**, our results reveal that the immune class is correlated with significantly lower copy number deletion, higher TILs abundance, higher TMB, higher PD-L1, but not neoantigens. The specific mutant genes in the immunophenotypes are diverse.

**Figure 5.**
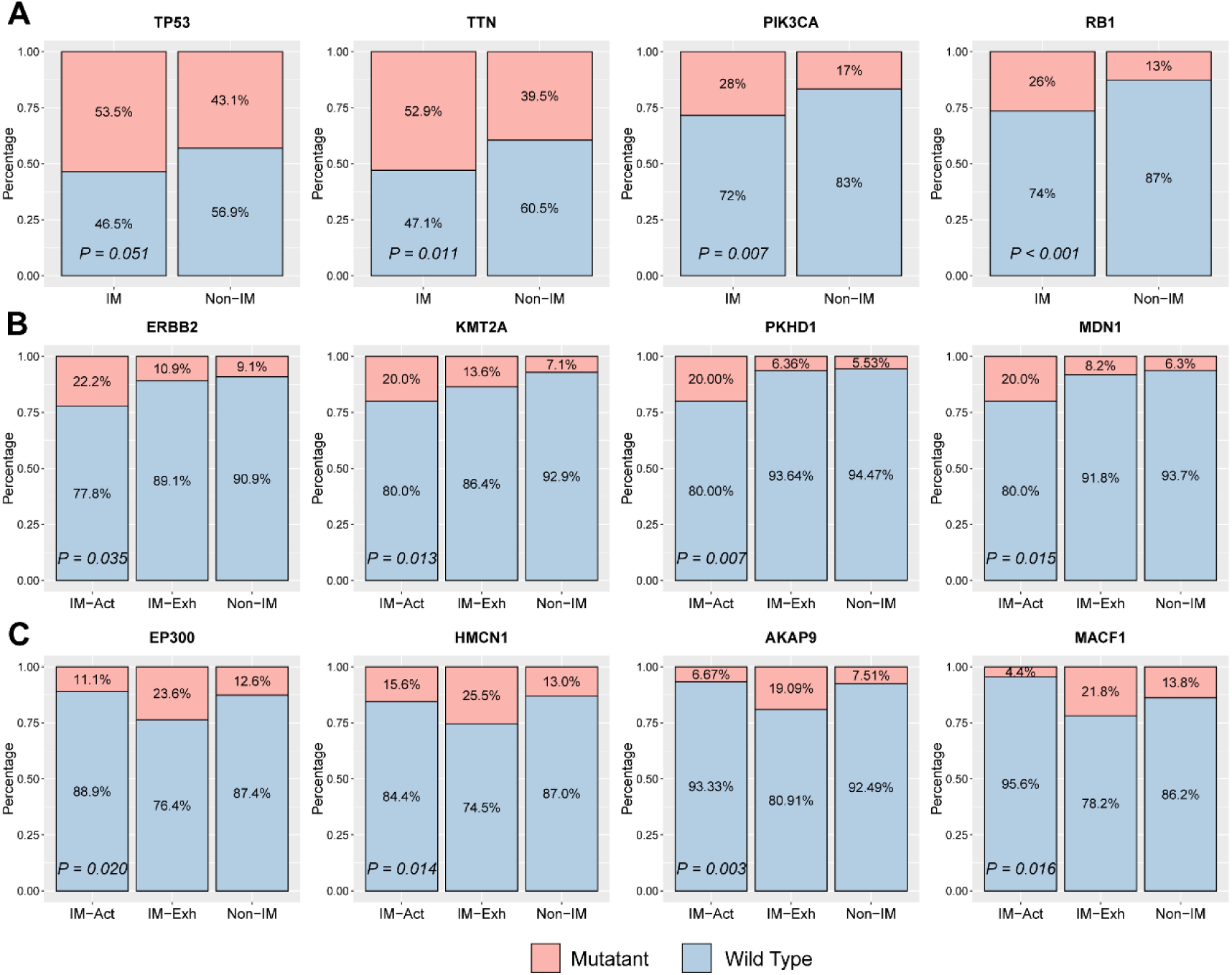
The specific mutant genes of non-immune class, immune activated subgroup, and immune exhausted subgroup. (A) TP53, TTN, PIK3CA and RB1 are the specific mutant genes in immune class compared with non-immune class; (B) ERBB2, KMT2A, PKHD1 and MDN1 are the specific mutant genes in immune activated subgroup; (C) EP300, HMCN1, AKAP9 and MACF1 are the specific mutant genes in immune exhausted subgroup. IM, immune class; Non-IM, non-immune class; IM-Act, immune activated subgroup; IM-Exh, immune exhausted subgroup.

### Reappearing the three immunophenotypes in 19 external cohorts

19 external cohorts with the mRNA expression profile were collected to reappear the three immunophenotypes defined by the NMF algorithm microdissected and activated stroma signature (**Figure 1, Table S1**). The increased top 150 different expression genes (DEGs) between the immune and non-immune classes (**Table S5**) was chosen as the seed genes to reappear the immune subclasses in the external cohorts with the GenePattern module “NMFConsensus”, and then, the immune class divided to activated and exhausted subgroups by the and nearest template prediction (NTP) module.

In GSE32894 cohort, 60.7% (187/308) patients identified as the non-immune class, with the lower enrichment of immune associated signatures, as for the remaining 121 patients, compared with the signatures of stromal enrichment, 42 patients belong to the immune activated subgroup, and 79 belong to the immune exhausted subgroup. Patients in the immune exhausted subgroup shown a high enrichment score of TITR, MDSC, WNT/TGFβ, TGFβ-1 activated and C-ECM signatures (all, *P <* 0.01, **Figure 6**).

**Figure 6.**
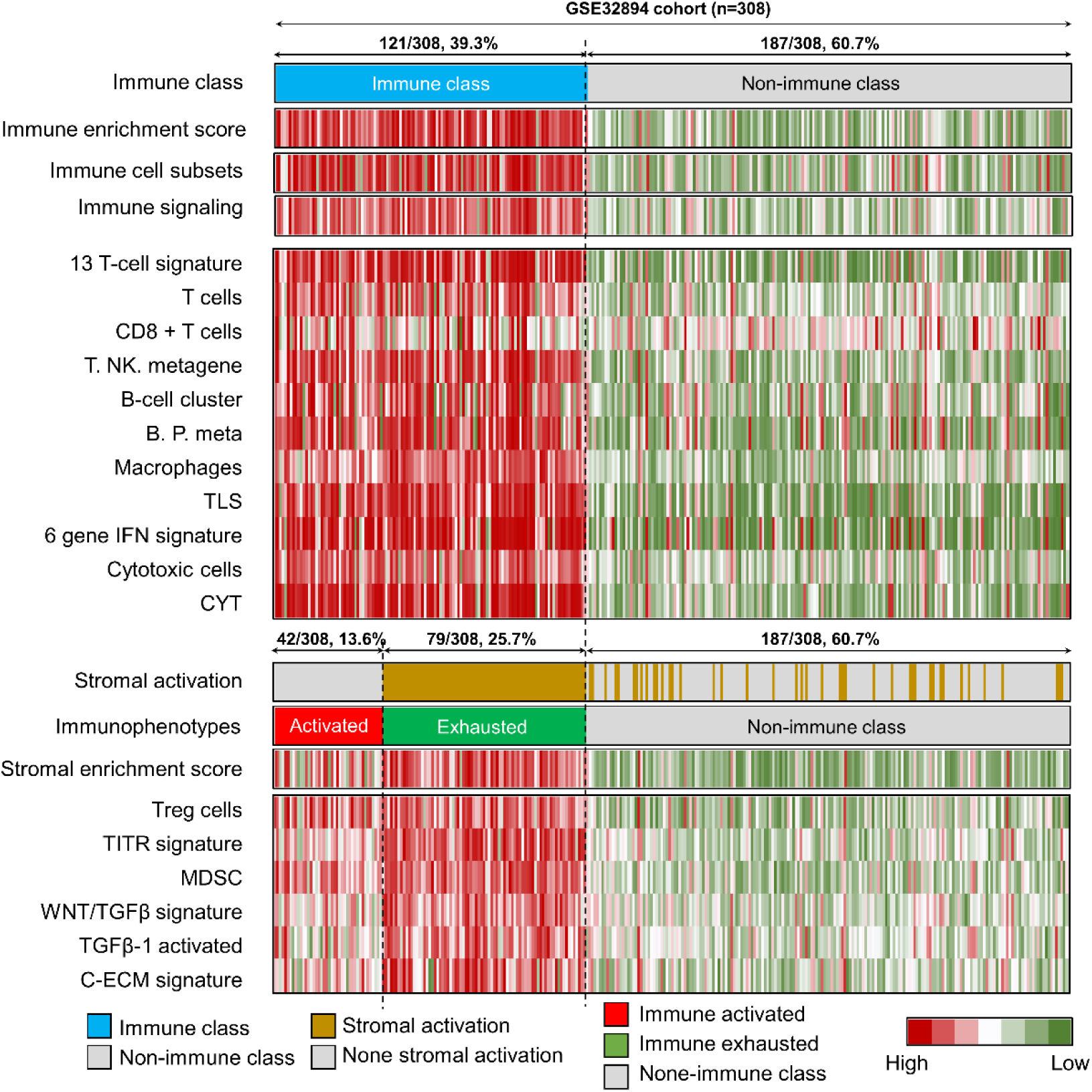
Reappearing the diverse immune characteristics of three immunophenotypes in GSE32894 cohort. CYT, cytolytic activity score; TITR, tumor-infiltrating Tregs; MDSC, myeloid-derived suppressor cell; TLS, tertiary lymphoid structure; C-ECM, cancer-associated extracellular matrix.

In the other 3287 bladder cancer patients from 18 cohorts, we also reappeared the three immunophenotypes, the results displayed in **Figure S4** to **Figure 21**. In these cohorts, the distribution of immune activated subgroups ranged from 11.3% to 30.9%, while the proportion of immune exhausted subgroups ranged from 17.1% to 40.8%. We also observed the increased scores of immune enrichment signature and immune signaling signature in the 18 validation cohorts, as well as the other immunocytes signatures. As expected, the subgroup of immune exhausted shown an increased enrichment score of Treg cells, TITR, MDSC, WNT/TGFβ, and C-ECM signatures. Taken together, combined results from **Table S5, Figure 6, and Figure S4 to S21**, our results suggest that the NMF and NTP algorithms could stably and precisely divide bladder patients into immune activated, immune exhausted and non-immune phenotypes. The specific immune characteristics could reappearing in any bladder cancer patients cohort.

### Immune activated subgroup shows favorable prognosis and benefits more from anti-PD-1 therapy

We collected the overall survival status and time form the TCGA-BLCA, GSE32894, GSE13507 and E-MTAB-1803 cohorts. The prognosis of patients in the three immunophenotypes are dramatically difference. In TCGA-BLCA cohort, we observed the best OS outcome in immune activated subgroup among patients older than 70 years old, while the survival plots of immune exhausted subgroup and non-immune class mixed (**Figure 7A**, *P =* 0.45). What’s more, the prognosis obviously distinguished between the three immunophenotypes in GSE32894 cohort (**Figure 7A**, *P <* 0.001). Patients in the immune activated subgroup all alive at the end of follow-up, patients belong to the non-immune class shown a low rate of death, 5.96% (9/151), while about one thirds patients (16/47) in the immune exhausted subgroup met the death end. The similar tendency of better prognosis in immune activated subgroup, worse prognosis in immune exhausted subgroup was also observed in E-MTAB-1803 cohort and GSE13507 cohort (**Figure 7A**).

**Figure 7.**
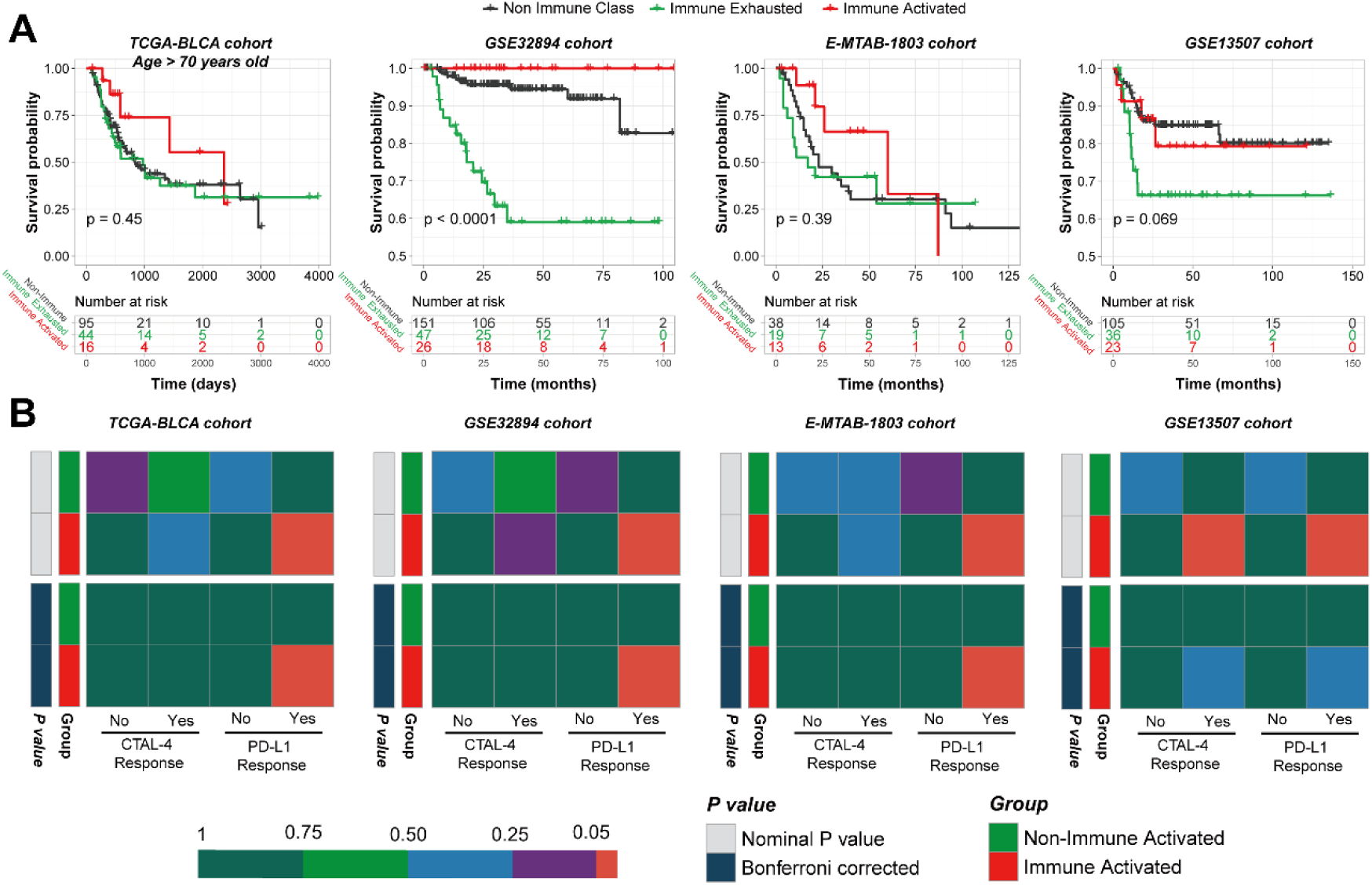
Immunophenotypes indicate separated overall survival outcome and response to immunotherapy for bladder cancer patients. (A) Different overall survival outcome in three immunophenotypes among patients high than 70 years old in TCGA-BLCA cohort, GSE32894 cohort, E-MTAB-1803 cohort, and GSE13507 cohort; (B) Subclass mapping analysis manifested that patients with immune activated subtype were more likely to respond to anti–PD-1 treatment.

Subsequently, we predict the potential response to anti-PD-1 and CTAL4 therapy of the patients in difference immunophenotypes. The module of SubMap in GenePattern was employed to compare the similarity of gene expression profile between the immunophenotypes and responders of anti-CTLA-4 or anti-PD-1 in the metastatic melanoma immunotherapy cohort. We successfully generated the results that patients in the immune activated subgroups could benefit more from the treatment of anti-PD-1 therapy but not anti-CTLA-4 therapy in TCGA-BLCA, GSE32894, and E-MTAB-1803 cohorts (Bonferroni-corrected *P <* 0.05, **Figure 7B**). Taken together of the results from **Figure 7**, we generated the conclusion of that patients in the immune activated subgroup have the longest average overall survival and could benefit more from the anti-PD-1 therapy.

### Correlate three immunophenotypes with proposed molecular subtypes

We also sought to integrate the immunophenotypes with the prior established immune molecular features. Thorsson *et al*.^35^ generated a six-subtype immune molecular feature, including wound healing (C1), IFN-γ dominant (C2), inflammatory (C3), lymphocyte depleted (C4), immunologically quiet (C5), and TGF-β dominant (C6). We found that most patients in the immune activated subgroup (75.00%) belong to the IFN-γ group, which associated with a strong CD8 signal and a high proliferation rate, and about 68.22% patients in the immune exhausted subgroup also belong to the IFN-γ group (**Figure 8A**).

**Figure 8.**
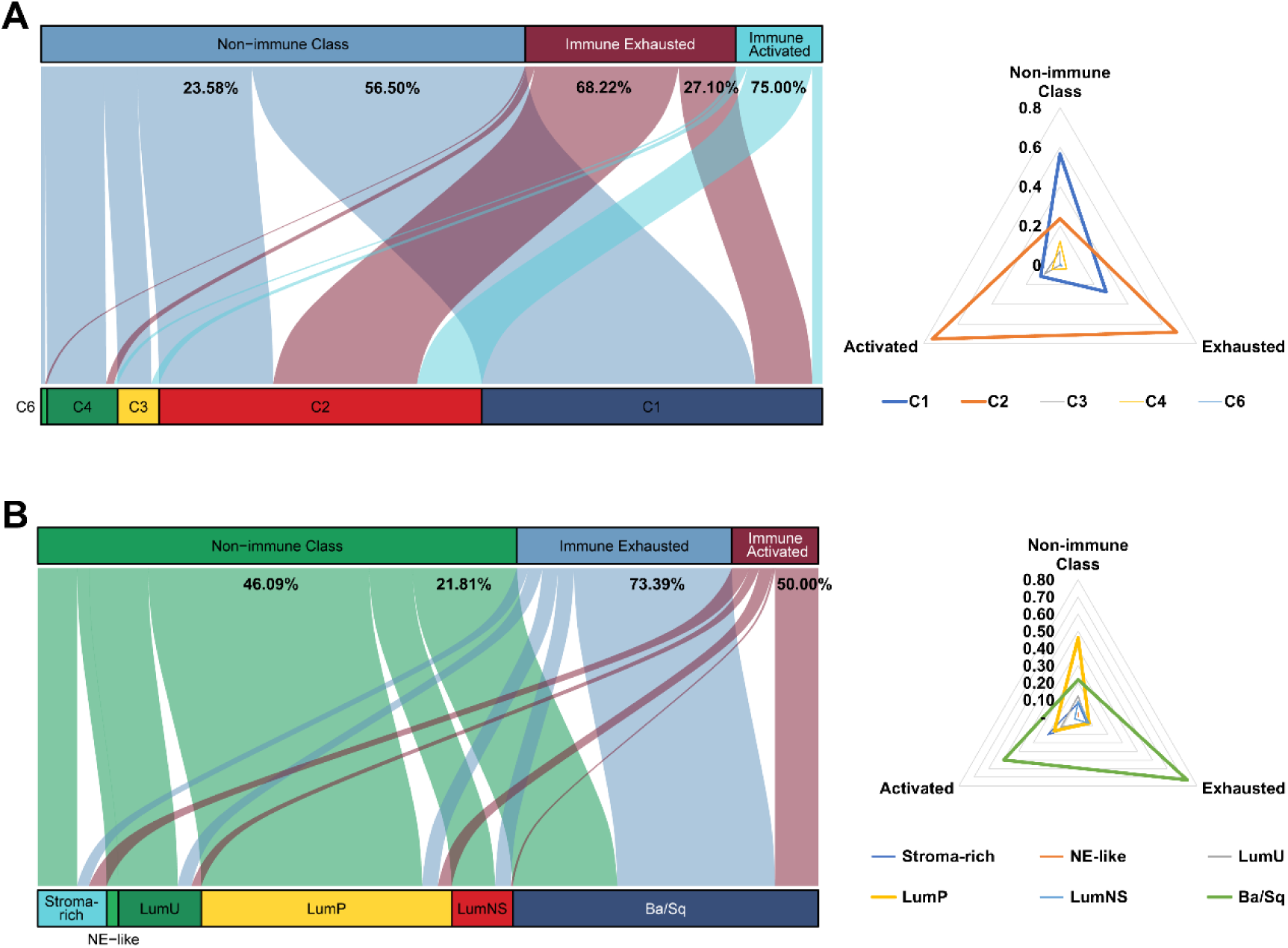
Correlate the three immunophenotypes with proposed molecular subtypes. (A) Association with Thorsson et al. generated pan-cancer six immune molecular features; (B) Association with Kamoun et al. identified the consensus set of six molecular classes.

Kamoun et al.^36^ identified a consensus set of six molecular classes: luminal papillary (24%), luminal nonspecified (8%), luminal unstable (15%), stroma-rich (15%), basal/squamous (35%), and neuroendocrine-like (3%). In the current study, we revealed that most immune exhausted patients and half of immune activated patients belong to the basal/squamous classes (**Figure 8B**), which with the most frequently mutated genes of TP53, consistent with what we generated previously (**Figure 5A**). Unquestionably, the Ba/sq subclass associated with the poor prognosis, and patients in the immune exhausted subgroup also shown the poor prognosis.

## DISCUSSION

Bladder cancer is a heavy health burden all over the world, especially at Europe and Northern America^2^. The major challenge of bladder cancer clinical care is the short-term recurrence of NMIBC, as well as the shorten overall survival of MIBC patients, especially for those with distant metastases, with the 5-year survival rate less than 10%^7,37^. Iridates from the molecular side, bladder cancer is composed by a mass of heterogenetic characteristics, impacting by the gene mutation, gene copy number alteration, neoantigens, as well as the infiltration of immunocytes. Several teams established the molecular classifications among bladder cancer. Mo et al.^38^generated a tumor 18-gene signature in MIBC patients, which could reflect the urothelial differentiation and predict the clinical outcomes, basal and differentiated groups was named to the two group with high or low risk score, respectively. Damrauer *et al*.^39^ developed BASE47, a transcriptomic classifier using 47 genes, to classify the MIBC tumor into luminal-like or basal-like subtype. Robertson et al.^40^ performed a Bayesian NMF with consensus hierarchical clustering in 408 MIBC tumors from TCGA and found five expression subtypes, including three luminal subtypes (named Luminal-papillary, Luminal-Infiltrated and Luminal), Basal/Squamous subtype, and Neuronal subtype. However, most of the molecular classifiers only focused on the clinical outcomes, but not the tumor immune microenvironment. Therefore, our goal is to provide a comprehensive insight to the immune response of bladder cancer patients with diverse inner molecular features, and help to find the suitable patients undergo the precise immunotherapy.

NMF algorithm is an unsupervised, parts-based learning paradigm, which could decompose a nonnegative matrix V into two nonnegative matrices, W and H, via a multiplicative updates algorithm^41^. Similarly to principal components analysis (PCA) or independent component analysis (ICA), the objective of NMF is to explain the observed data using a limited number of basic components, which could reflect the original data as accurately as possible^42^. NMF was applied to reveal the biomarkers, classify the tumor subtypes and predict the prognosis of tumors recent days^43-45^. As for the enrolled 4,003 bladder cancer patients. We investigated a robust classification of three immunophenotypes based on the NMF algorithm. Firstly, we identified the immune activated subgroup, immune exhausted subgroup and non-immune class in the 408 bladder cancer patients from TCGA-BLCA cohort. The 150 exemplar genes from immune module represented the immune feature in bladder cancer patients and further divide the whole cohort to immune and non-immune classes. The other 150 DEGs among the immune and non-immune classes was extracted as the input profile for the validation of the classification in external 19 cohorts. As to the distinguish of immune activated and immune exhausted subgroup, a stromal activation signature was conducted by NTP algorithm. The features of these three immunophenotypes was illuminated by several verified signatures of immunocytes or immune signaling pathways. Patients in the immune classes shown the highly enriched signatures of T cell, B cell, IFN and CYT^46-48^, while the exhausted subgroup also shown an increased signature of TITR, WNT/TGF-β, TGF-β1 activated, and C-ECM signatures^49-51^, but not the immune activated subgroup. Totally, based on our results, we revealed that there are only about 11% to 30.9% bladder cancer patients belong to the immune activated subgroup, which might response to the immunotherapy.

Clinical outcome is an important factor we focused about the newly defined immunophenotypes. With the clinical information of TCGA-BLCA, GSE32894, E-MTAB-1803 and GSE13507 cohorts, we generated the results that patients belong to the immune activated subgroup contain the best overall survival, while the immune exhausted subgroup shown the worst clinical outcome of a shorten overall survival time. Immune exhausted, mostly focused on the exhausted of T cell, reflected by the altered inflammatory and tissue microenvironments, lymphocyte, as well as the inhibitory signals from cytokines^52^. These alternations in the TIME could lead to the escape of the immune recognition by blocking of the immune checkpoints, and related with the unfavorable overall survival for patients^53^. We predicted the potential response to immunotherapy of the bladder cancer patients by compared the mRNA expression profile with melanoma samples receiving anti-CTLA-4 or anti-PD-1 checkpoint therapy. As expected, patients in the immune activated subgroup could benefit from the treatment of an-PD-1 therapy, but not the non-immune activated subgroup, which combined the immune exhausted and non-immune classes.

To further understanding the molecular diverse among there three immunophenotypes, we compared the CNA, TMB, and gene mutations. Recent studies report the association of CNA with the increased immune infiltration and the outcome of immune checkpoint blockade therapy^54,55^. Patients in the immune class shown a lower CNA burden in gene deletion among arm- and focal-level. The association was double checked by that the deletion copy number of PD-1, PD-L1 and CTLA4 is positively with the decrease infiltration level of immunocytes. Tripathi et al.^56^ found that antigen presentation through MHC class I pathway is suppressed in tumors with high chromosomal instability, also known as the high CNA, which acts as a pivotal role in the immune evasion. In addition, Lu et al.^57^ revealed that patients treated with immune-checkpoint-blockade therapy could get a durable clinical benefit and better survival, if the contains the lower burden of CNA. Gene mutation is another key component we focused among the three immunophenotypes. We extracted the specific mutant genes for each subgroup. The proportion of mutant TP53, TTN, PIC3CA and RB1 is higher in immune class than non-immune class. Nusrat et al.^58^ reported that colorectal cancer patients with the mutant PIK3CA have a higher median density of CD3+ and CD8+ cells, as well as a high rate of clinical benefit form immunotherapy (50% vs. 8.6%). What’s more, we observed the high rate of ERBB2 mutation in immune activated subgroup. ERBB2 amplification or overexpression was a biomarker of anti-ERBB2 target therapy in breast cancer, the activated ERBB2 oncogene regulates recruitment and activation of tumor infiltrating immune cells and trastuzumab activity by inducing CCL2 and PD-1 ligands^59^. The V659E mutation of ERBB2 gene was also reported associated with the altering sensitivity of afatinib and lapatinib treatment in in vitro^60,61^. The mutation proportion of EP300 is highest in immune exhausted subgroup. Recent research concerned the importance of CBP/EP300 in regulatory T cells (Treg), because conditional deletion of either EP300 or CBP in mouse Tregs led to impaired Treg suppressive function^62^. Intratumoral Tregs dampen effector T cell responses to tumor antigens, engendering an immunosuppressive microenvironment, and linked with a poor prognosis in tumors^63^.

We defined and validated a novel classifier among the 4003 bladder cancer patients, to separate the bladder cancer patients to immune activated, immune suppressed and non-immune subgroups. Patients in the immune activated subgroup could benefit more from the single treatment of anti-PD-1 immunotherapy; As to the immune exhausted subgroup, ICB therapy plus TGF-β inhibitor or EP300 inhibitor might be more effectiveness. In summary, our novel classifier provide illumination for the enhancing immunotherapy of bladder cancer patients.

## Data availability statement

All data used in this work can be acquired from the GDC portal (https://portal.gdc.cancer.gov/), Gene-Expression Omni-bus (GEO; https://www.ncbi.nlm.nih.gov/geo/), and ArrayExpress (https://www.ebi.ac.uk/arrayexpress/).

## Acknowledgements

This work was supported by the National Natural Science Foundation of China [grant number: 81802827, 81630019, 31701162]; Scientific Research Foundation of the Institute for Translational Medicine of Anhui Province [grant number: 2017ZHYX02]; The Natural Science Foundation of Guangdong Province, China [grant number: 2017A030313800]; The Key Project of Provincial Natural Science Research Project of Anhui Colleges [grant number: KJ2019A0278]; Supporting Project for Distinguished Young Scholar of Anhui Colleges [grant number: gxyqZD2019018]; 2017 Anhui Province special program for guiding local science and technology development by the central government [grant number: 2017070802D148].

## Ethics approval

The patient data in this work acquired from the publicly available datasets whose informed consent of patients were complete. For the AHMU-PC cohort, the research contents and research programs were reviewed and approved by the Ethics Committee of the First Affiliated Hospital of Anhui Medical University (PJ-2019-09-11), patient consent for the retrospective cohorts was waived.

## Authors’ contributions

Conception and Design: Jialin Meng, Xiaofan Lu, Fangrong Yan and Chaozhao Liang. Collection and Assembly of Data: Yujie Zhou, Xiaofan Lu, Meng Zhang, Yinan Du, Jun Zhou. Data Analysis and Interpretation: Jialin Meng, Yujie Zhou, Xiaofan Lu, Zongyao Hao. Manuscript Writing: Jialin Meng, Xiaofan Lu, Yujie Zhou, Meng Zhang. Final Approval of Manuscript: All the authors.

## Figure Legends

**Figure S1. GSEA results showing the activated signaling pathways in the immune class**. NES, normalized enrichment score; FDR, false discovery rate; FDR less than 0.05 indicates statistical significance.

**Figure S2. Stromal representative signatures and markers in immune activated and exhausted subgroups**. ****, *P <* 0.0001; ***, *P <* 0.001; **, *P <* 0.01; *, *P <* 0.05; ns, no significance.

**Figure S3. The association between copy number variation of immune checkpoints and immunocyte infiltration**. ***, *P <* 0.001; **, *P <* 0.01; *, *P <* 0.05.

**Figure S4. Successful validation of the immunophenotypes among the E-MTAB-4321 cohort**. CYT, cytolytic activity score; TITR, tumor-infiltrating Tregs; MDSC, myeloid-derived suppressor cell; TLS, tertiary lymphoid structure; C-ECM, cancer-associated extracellular matrix.

**Figure S5. Successful validation of the immunophenotypes among the IMvigor210 cohort**. CYT, cytolytic activity score; TITR, tumor-infiltrating Tregs; MDSC, myeloid-derived suppressor cell; TLS, tertiary lymphoid structure; C-ECM, cancer-associated extracellular matrix.

**Figure S6. Successful validation of the immunophenotypes among the GSE83586 cohort**. CYT, cytolytic activity score; TITR, tumor-infiltrating Tregs; MDSC, myeloid-derived suppressor cell; TLS, tertiary lymphoid structure; C-ECM, cancer-associated extracellular matrix.

**Figure S7. Successful validation of the immunophenotypes among the GSE87304 cohort**. CYT, cytolytic activity score; TITR, tumor-infiltrating Tregs; MDSC, myeloid-derived suppressor cell; TLS, tertiary lymphoid structure; C-ECM, cancer-associated extracellular matrix.

**Figure S8. Successful validation of the immunophenotypes among the GSE128702 cohort**. CYT, cytolytic activity score; TITR, tumor-infiltrating Tregs; MDSC, myeloid-derived suppressor cell; TLS, tertiary lymphoid structure; C-ECM, cancer-associated extracellular matrix.

**Figure S9. Successful validation of the immunophenotypes among the GSE13507 cohort**. CYT, cytolytic activity score; TITR, tumor-infiltrating Tregs; MDSC, myeloid-derived suppressor cell; TLS, tertiary lymphoid structure; C-ECM, cancer-associated extracellular matrix.

**Figure S10. Successful validation of the immunophenotypes among the GSE120736 cohort**. CYT, cytolytic activity score; TITR, tumor-infiltrating Tregs; MDSC, myeloid-derived suppressor cell; TLS, tertiary lymphoid structure; C-ECM, cancer-associated extracellular matrix.

**Figure S11. Successful validation of the immunophenotypes among the GSE39016 cohort**. CYT, cytolytic activity score; TITR, tumor-infiltrating Tregs; MDSC, myeloid-derived suppressor cell; TLS, tertiary lymphoid structure; C-ECM, cancer-associated extracellular matrix.

**Figure S12. Successful validation of the immunophenotypes among the GSE128701 cohort**. CYT, cytolytic activity score; TITR, tumor-infiltrating Tregs; MDSC, myeloid-derived suppressor cell; TLS, tertiary lymphoid structure; C-ECM, cancer-associated extracellular matrix.

**Figure S13. Successful validation of the immunophenotypes among the GSE124035 cohort**. CYT, cytolytic activity score; TITR, tumor-infiltrating Tregs; MDSC, myeloid-derived suppressor cell; TLS, tertiary lymphoid structure; C-ECM, cancer-associated extracellular matrix.

**Figure S14. Successful validation of the immunophenotypes among the GSE86411 cohort**. CYT, cytolytic activity score; TITR, tumor-infiltrating Tregs; MDSC, myeloid-derived suppressor cell; TLS, tertiary lymphoid structure; C-ECM, cancer-associated extracellular matrix.

**Figure S15. Successful validation of the immunophenotypes among the GSE48276 cohort**. CYT, cytolytic activity score; TITR, tumor-infiltrating Tregs; MDSC, myeloid-derived suppressor cell; TLS, tertiary lymphoid structure; C-ECM, cancer-associated extracellular matrix.

**Figure S16. Successful validation of the immunophenotypes among the GSE31684 cohort**. CYT, cytolytic activity score; TITR, tumor-infiltrating Tregs; MDSC, myeloid-derived suppressor cell; TLS, tertiary lymphoid structure; C-ECM, cancer-associated extracellular matrix.

**Figure S17. Successful validation of the immunophenotypes among the GSE134292 cohort**. CYT, cytolytic activity score; TITR, tumor-infiltrating Tregs; MDSC, myeloid-derived suppressor cell; TLS, tertiary lymphoid structure; C-ECM, cancer-associated extracellular matrix.

**Figure S18. Successful validation of the immunophenotypes among the GSE93257 cohort**. CYT, cytolytic activity score; TITR, tumor-infiltrating Tregs; MDSC, myeloid-derived suppressor cell; TLS, tertiary lymphoid structure; C-ECM, cancer-associated extracellular matrix.

**Figure S19. Successful validation of the immunophenotypes among the E-MTAB-1803 cohort**. CYT, cytolytic activity score; TITR, tumor-infiltrating Tregs; MDSC, myeloid-derived suppressor cell; TLS, tertiary lymphoid structure; C-ECM, cancer-associated extracellular matrix.

**Figure S20. Successful validation of the immunophenotypes among the GSE69795 cohort**. CYT, cytolytic activity score; TITR, tumor-infiltrating Tregs; MDSC, myeloid-derived suppressor cell; TLS, tertiary lymphoid structure; C-ECM, cancer-associated extracellular matrix.

**Figure S21. Successful validation of the immunophenotypes among the GSE129871 cohort**. CYT, cytolytic activity score; TITR, tumor-infiltrating Tregs; MDSC, myeloid-derived suppressor cell; TLS, tertiary lymphoid structure; C-ECM, cancer-associated extracellular matrix.

